# AI boosted molecular MRI for apoptosis detection in oncolytic virotherapy

**DOI:** 10.1101/2020.03.05.977793

**Authors:** Or Perlman, Hirotaka Ito, Kai Herz, Naoyuki Shono, Hiroshi Nakashima, Moritz Zaiss, E. Antonio Chiocca, Ouri Cohen, Matthew S. Rosen, Christian T. Farrar

**Affiliations:** Athinoula A. Martinos Center for Biomedical Imaging, Department of Radiology, Massachusetts General Hospital and Harvard Medical School, Charlestown, MA, USA; Brigham and Womens Hospital and Harvard Medical School, Boston, MA, USA; Magnetic Resonance Center, Max Planck Institute for Biological Cybernetics, Tübingen, Germany; IMPRS for Cognitive and Systems Neuroscience, University of Tübingen, Tübingen, Germany; Department of Neuroradiology, University Clinic Erlangen, Erlangen, Germany; Memorial Sloan Kettering Cancer Center, New York, New York; Department of Physics, Harvard University, Cambridge, MA, USA

## Abstract

Oncolytic virotherapy is a promising treatment for high mortality cancers^1^. Non-invasive imaging of the underlying molecular processes is an essential tool for therapy optimization and assessment of viral spread, innate immunity, and therapeutic response^2, 3^. However, previous methods for imaging oncolytic viruses did not correlate with late viral activity^4^ or had poor sensitivity and specificity^5^. Similarly, methods developed to image treatment response, such as apoptosis, proved to be slow, nonspecific, or require the use of radioactive or metal-based contrast agents^6–8^. To date, no method has been widely adopted for clinical use. We describe here a new method for fast magnetic resonance molecular imaging with quantitative proton chemical-exchange specificity to monitor oncolytic virotherapy treatment response. A deep neural network enabled the computation of quantitative biomarker maps of protein and lipid/macromolecule concentrations as well as intracellular pH in a glioblastoma multiforme mouse brain tumor model. Early detection of apoptotic response to oncolytic virotherapy, characterized by decreased cytosolic pH and protein synthesis, was observed in agreement with histology. Clinical translation was demonstrated in a normal human subject, yielding molecular parameters in good agreement with literature values^9^. The developed method is directly applicable to a wide range of pathologies, including stroke^10^, cancer^11–13^, and neurological disorders^14, 15^.

---

The highly invasive nature of many cancer types and the toxicity of most systemic chemotherapies represent significant challenges for cancer therapies and limit their effectiveness. An especially promising therapeutic approach for overcoming these challenges is the use of oncolytic viruses that selectively kill only cancer cells while sparing the surrounding normal cells^16^. Oncolytic viruses can generate progeny on-site that spread throughout the tumor and reach distal malignant cells, thus, representing an ideal strategy for treating invasive cancers such as Glioblastoma^17^. Oncolytic viruses can also be armed to express anticancer genes and provide targeted delivery of therapeutics^18, 19^. In addition, oncolytic viruses can elicit a strong immune response against virally-infected tumor cells and were recently FDA approved for melanoma treatment^20^. Non-invasive methods to image oncolytic viruses are essential for quantifying virus titer and achieving the full potential of this biological therapeutic^4^. In-vivo molecular information provided during the course of therapy could provide detailed insights into both the tumor and host response and help optimize and expand the current scientific horizons of virotherapy.

Chemical exchange saturation transfer (CEST) magnetic resonance imaging (MRI) has previously been shown to be sensitive to changes in tumor pathology^12^ and could provide such a non-invasive imaging tool. CEST is a molecular imaging technique that uses radio-frequency (RF) pulses to saturate the magnetization of exchangeable protons on a variety of molecules, including proteins and metabolites^21^. This generates a detectable MRI signal change due to fast chemical exchange with the abundant water pool. The CEST contrast depends on the chemical exchange rate, which is pH sensitive, and the volume-fraction of the exchangeable proton pool, which is sensitive to protein and metabolite concentrations. The sensitivity of CEST MRI to pH and protein/metabolite concentrations has proven to be a potent tool for imaging a wide range of pathologies^22^. However, the clinical translation of CEST-MRI methods has been hindered by the qualitative nature of the image contrast and the typically long image acquisition times required.

Here, we report the design of a deep learning based CEST fingerprinting method for quantitative and rapid molecular imaging of oncolytic virotherapy (OV) treatment response without the need for any exogenous contrast agents. The proposed technique is based on selective magnetic labeling of exchangeable amide protons of endogenous proteins as well as exchangeable protons of lipids and macromolecules (Fig. 1a) using a pseudo-random and fast (3.5 min) RF saturation pulse scheme (Fig. 1b), which encodes the molecular properties into unique MR-fingerprints^23^. Next, the acquired signals are rapidly decoded (94 ms) into four fully quantitative chemical exchange and proton concentration molecular maps, using a series of deep neural networks (Fig. 1c), trained with a dictionary of simulated MR-fingerprints. The dictionaries are simulated for a range of semi-solid and amide chemical exchange parameter values as well as water T_1_ and T_2_ relaxation times and B_0_ magnetic field homogeneity values. The incorporation of the artificial intelligence (AI)-based reconstruction within the system architecture provides the ability to overcome the highly multi-dimensional nature of this fingerprint matching problem and successfully decouple and quantify the different molecular properties. Moreover, the reconstruction time is five orders of magnitude faster than traditional MR-fingerprinting reconstruction, contributing to the clinical translation potential.

**Figure 1:**
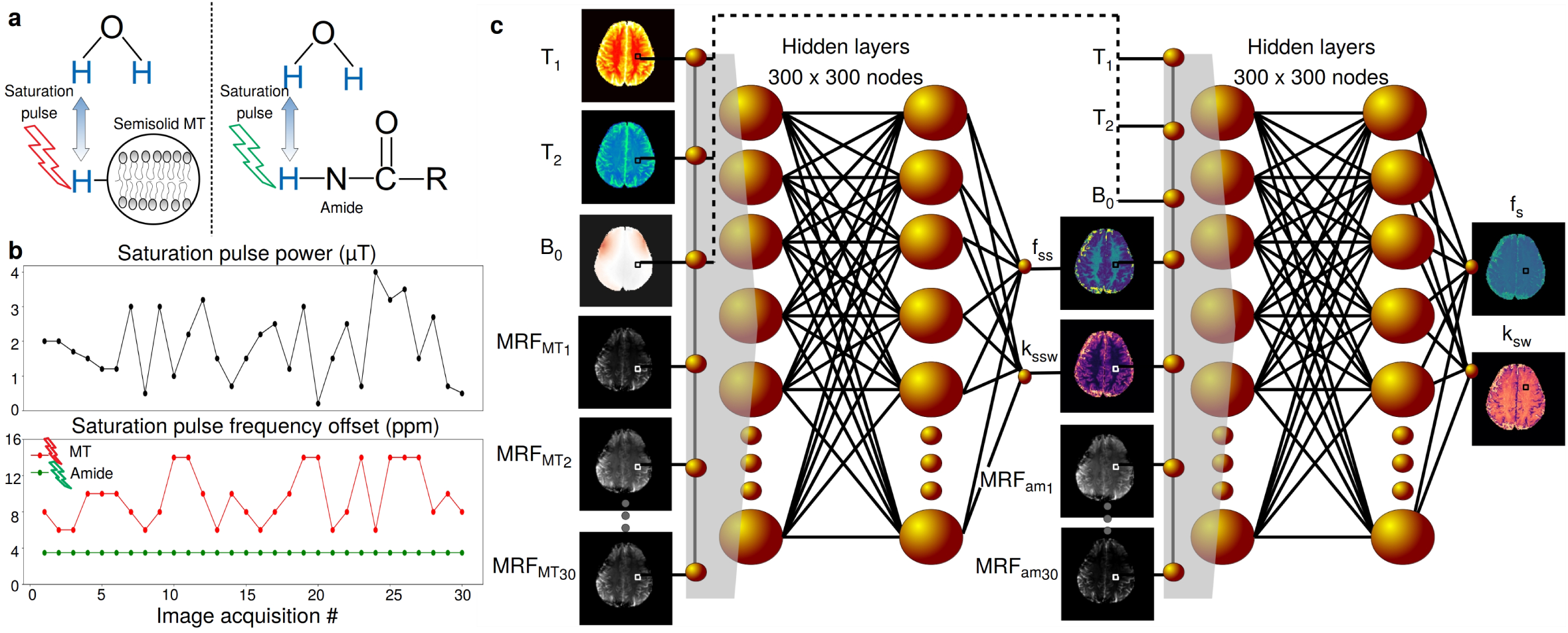
Schematic representation of AI-boosted molecular MRI pipeline. a. The molecular information of the compounds of interest (semisolid MT and amide) is encoded into unique MR-fingerprints, using a series of saturation pulses (described in b). This results in two sets of raw molecular-feature-embedded images (MRF_*MT*_ and MRF_*am*_). c. Quantitative image decoding. The encoded image-sets as well as quantitative water-pool and field homogeneity maps (T_1_, T_2_, B_0_) are input sequentially into two deep reconstruction neural networks, ultimately yielding quantitative molecular maps, depicting the proton exchange rate and volume fraction for the semisolid and amide pools (k_*ssw*_, f_*ss*_, k_*sw*_, and f_*s*_, respectively).

The approach was evaluated in mice undergoing virotherapy treatment. The resulting quantitative maps allowed for the early detection of apoptosis induced by oncolytic virotherapy. The method was translated to clinical MRI scanners and used to image a healthy human subject, providing quantitative molecular maps in good agreement with the literature.

## Results

The suitability of the AI-boosted molecular MRI method for monitoring oncolytic virotherapy treatment response was evaluated in a longitudinal animal study. A preclinical orthotopic mouse model of a U87ΔEGFR human glioblastoma was used (n=16, 25% served as control). Imaging was performed at baseline (8-11 days post tumor implantation) as well as 48 and 72 hours post-OV treatments.

The quantitative exchange parameter-maps obtained for a representative virotherapy treated mouse can be seen in Fig. 2a-c, and the quantitative analysis for all OV-treated mice is presented in Fig. 2d. For all examined molecular parameters, the null hypothesis, claiming that the tumor, contralateral, and apoptotic region at all time-points are from the same distribution and have equal means was rejected by one-way analysis of variance (ANOVA) (F(7, 64)=45.87, 18.24, 95.09, and 13.14 for the amide proton volume fraction and exchange rate, and the semi-solid proton volume fraction and exchange rate, respectively, P<0.0001 in all cases). All group comparison analyses were performed using Tukeys multiple comparisons test (see Methods section).

**Figure 2:**
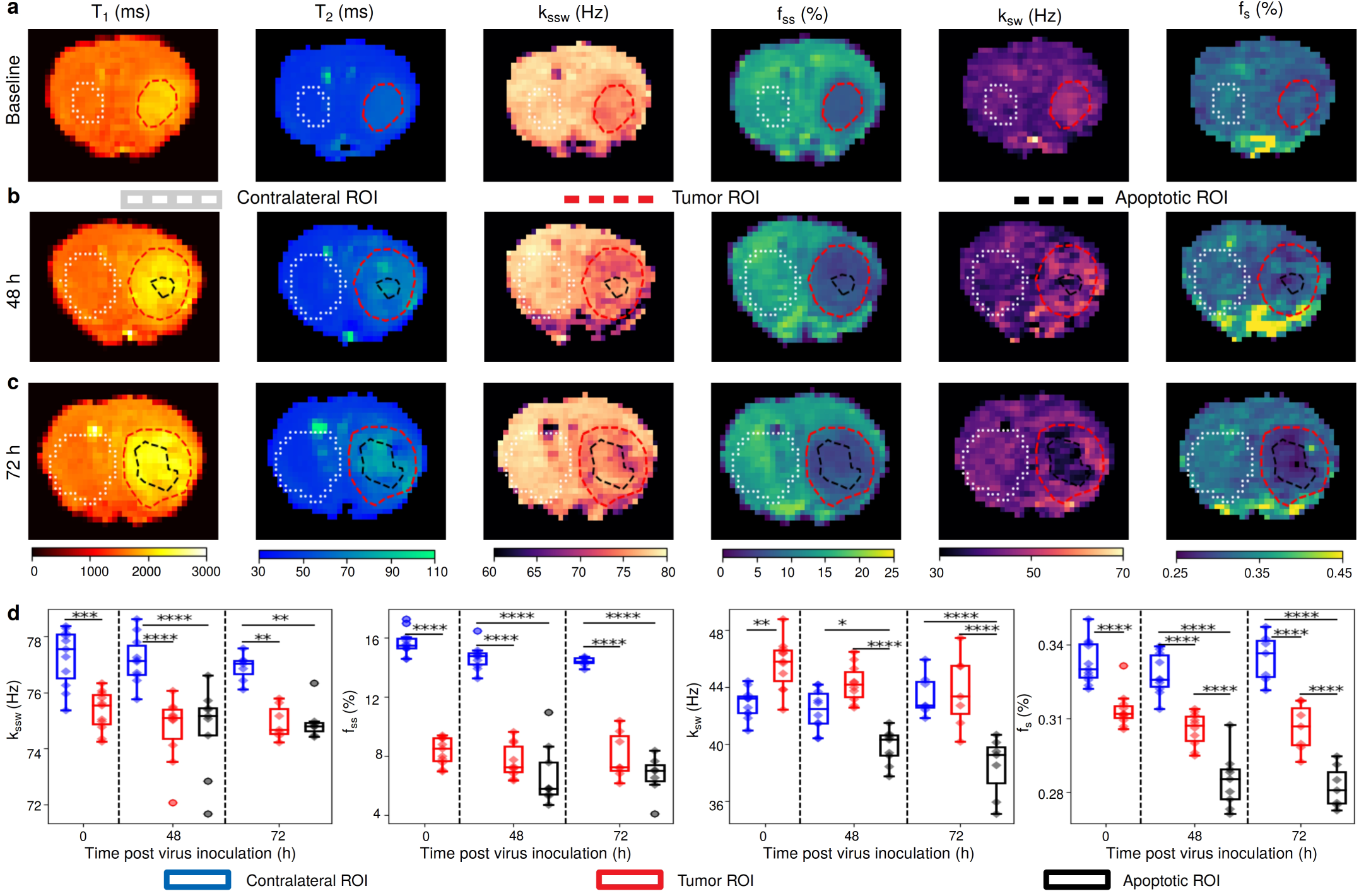
Quantitative molecular images of a representative oncolytic virotherapy (OV) treated mouse. a. Before inoculation the tumor semi-solid (f_*ss*_) and amide (f_*s*_) proton concentrations were decreased, consistent with increased edema. The tumor amide proton exchange-rate (k_*sw*_) was increased, indicative of increased intracellular pH. Forty-eight (b) and 72 (c) hours following OV, the tumor core presented significantly lower f_*s*_ and k_*sw*_ compared to the tumor rim and the contralateral region, indicative of apoptosis. d. Quantitative group comparison, demonstrating the statistical significance of the described phenomena using one-way analysis of variance (ANOVA) (F(7, 64)=45.87, 18.24, 95.09, and 13.14 for f_*s*_, k_*sw*_, f_*ss*_, and semi-solid exchange rate (k_*ssw*_), respectively, P<0.0001 in all cases) and Tukeys multiple comparison test, baseline (n=11 all regions), 48 hours (n=10 contralateral and tumor; n=9, apoptotic), and 72 hours (n=7, all regions) post inoculation. *P<0.05; **P<0.01; ***P<0.001; ****P<0.0001. The box plots describe the median, Tukey style whiskers, and outliers.

Prior to OV inoculation, the semi-solid (f_*ss*_) and amide proton (f_*s*_) volume fractions were both found to be significantly decreased in the tumor compared to the contralateral tissue (P<0.0001, n = 11), consistent with increased tumor edema decreasing the protein concentrations (Fig. 2a). In contrast, the tumor amide proton exchange-rate (k_*sw*_) was significantly increased (P<0.01, n = 11), indicative of increased intracellular pH in agreement with literature reports^24^. Forty-eight hours following OV (Fig. 2b), the core of the tumor presented significantly lower amide proton concentration compared to the contralateral (P<0.0001, n=10 contralateral, n=9 tumor core) and tumor rim regions (P<0.0001, n=10 tumor rim; n=9, tumor core). The same trend was observed at 72 hours post inoculation (P<0.0001, n=7). The amide proton exchange rate at 48 hours following OV was significantly lower in the tumor core compared to the contralateral (P=0.0197, n=10 contralateral, n=9 tumor core) and tumor rim regions (P<0.0001, n=10 tumor rim; n=9, tumor core). A similar trend was observed at 72 hours post inoculation (P<0.0001, n=7). The decrease in both amide proton exchange rate (which is sensitive to pH) and volume fraction (which is sensitive to protein concentration) suggests an apoptotic event in these areas as it is known to inhibit protein synthesis^25^ and decrease cytosolic pH^26^. Interestingly, the semi-solid exchange rate was significantly decreased in the tumor compared to the contralateral region at all time points (P<0.001, P<0.0001, and P<0.01 for baseline (n=11), 48h (n=10 contralateral and tumor; n=9, apoptotic), and 72h (n=7) post inoculation, respectively). This is attributed to the change in lipid composition of the tumor cell membranes compared to the healthy brain tissue^27^, which alters the base catalyzed exchange rate constant of the semi-solid protons^28^. Thus, the semi-solid proton exchange rate depends not only on pH, but also on the lipid/macromolecule composition, leading to a decreased exchange rate at baseline despite the increased pH. In contrast, for amide exchangeable protons from small mobile proteins with simple aqueous chemical environments, the base catalyzed exchange rate constant remains constant and the exchange rate is dependent only on the pH. The reproducibility of the proposed imaging method was confirmed by the lack of statistically significant differences in the parameter values of the contralateral region over-time (amide proton volume fraction: P>0.78 and proton exchange rate: P>0.86; semi-solid macro-molecules volume fraction: P>0.41 and exchange rate: P>0.49, minimal P-value is mentioned for baseline (n=11 all regions), 48 hours (n=10 contralateral and tumor; n=9, apoptotic), and 72 hours (n=7, all regions) post inoculation).

We next validated the MRI-based molecular findings using histology and immunohistochemistry (IHC). Formalin-fixed paraffin-embedded (PPFE) tissue sections were extracted from randomly chosen mice. A representative histology/IHC image-set and its comparison to the corresponding MR image-set can be seen in Fig. 3. HSV-1 antigens (indicating the viral biodistribution) were detected by IHC (Fig. 3e) and were located within the tumor boundaries (Fig. 3f), marked by the hematoxylin and eosin (HE) stained region. The HE stained region was in good agreement with the region of decreased MR semi-solid proton volume fraction (Fig. 3b). A well-defined region of IHC positive cleaved caspase 3 fragment, indicative of cell apoptosis, was observed (Fig. 3g) in good agreement with the region of reduced amide proton exchange rate (Fig. 3c). The Coomassie Blue stained image indicated that a reduction in protein concentration occurred at the tumor core (Fig. 3h), in a region similar to the apoptotic one (Fig. 3g) and in agreement with the region of decreased amide proton volume fraction (Fig. 3d).

**Figure 3:**
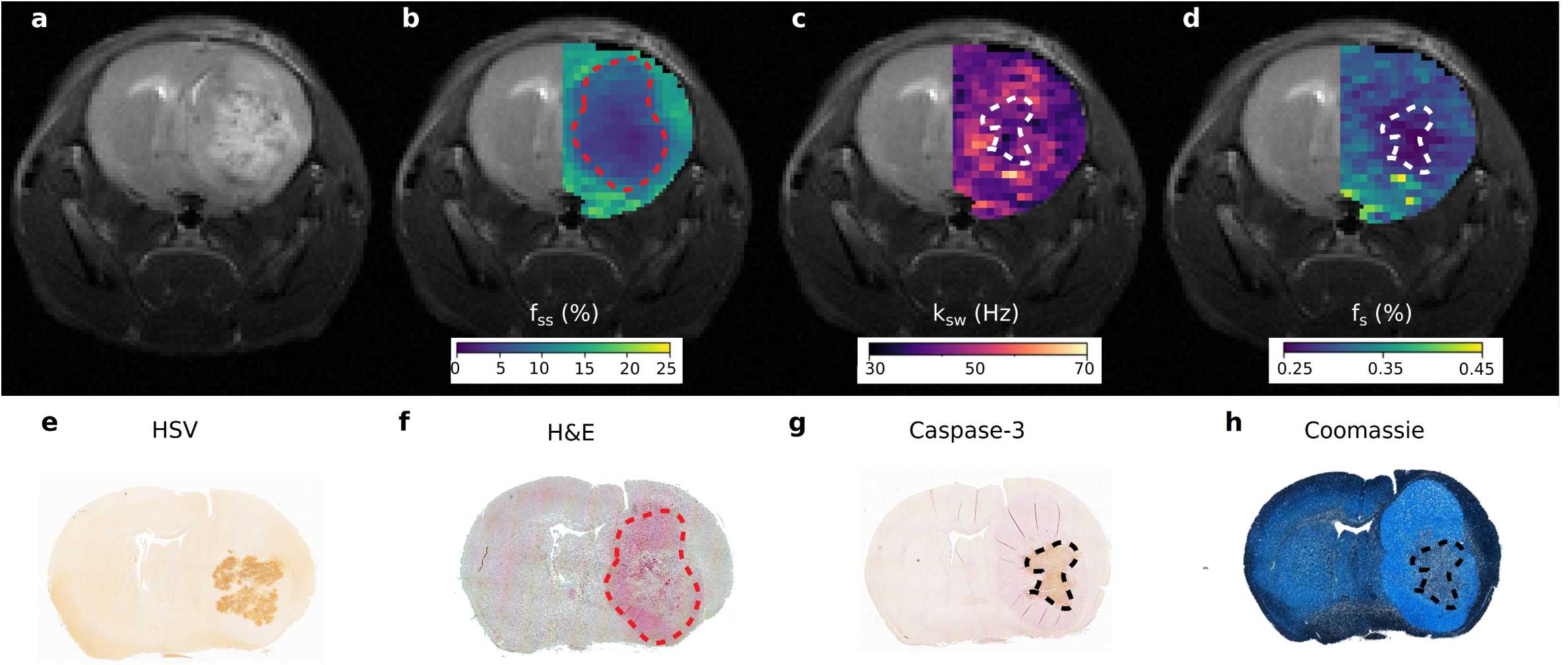
Histology validation. a. T_2_-weighted image of an OV-treated mouse, 72 hours post virus inoculation. b. Semisolid macro-molecules proton volume fraction (f_*ss*_) map, overlaid atop the T_2_-weighted image at the ipsilateral side. c. Similarly overlaid amide proton exchange-rate (k_*sw*_) and d. amide proton volume fraction (f_*s*_) maps. e. Immunohistochemistry image stained for Herpes Simplex Virus (HSV) presence (brown). f. HE stained image, demonstrating the tumor location (pink). g. Caspase-3 immunohistochemistry image, demonstrating the apoptotic tumor region (brown). h. Coomassie Blue stained image, demonstrating reduced protein concentration in the apoptotic tumor core. The dashed lines in images b-d, and f-h, generally depict the tumor (b, f) and apoptotic (c, d, g, h) regions borders, respectively.

The quantitative maps obtained for a representative non-virally treated control mouse and the quantitative analysis of the entire control group can be seen in Extended Data Fig. 1. The trends in the molecular exchange parameters at baseline were similar to that occurring for the OV-treated group (Fig. 2a), as expected. Namely, an apparent decrease in amide proton volume fraction, semi-solid proton volume fraction, and semi-solid proton exchange rate, at the tumor region, accompanied by a simultaneous increase in the tumor amide-proton exchange rate. We note that although the effect was statistically significant for the semi-solid exchange rate and volume fraction (one way ANOVA (F(5, 12)= 9.598, 100.8, P=0.0007, P<0.0001, respectively, and Tukey’s multiple comparisons test P<0.05, P<0.0001, respectively, n=3), it was not significant for the amide proton volume fraction and exchange rate (one way ANOVA (F(5, 12)=3.804, 0.8968, P=0.0269, P=0.5136, respectively, and Tukey’s multiple comparisons test P>0.05, n=3). As expected, at the later time points, no apoptotic region, manifested as a region of decreased amide exchange rate, was detected (Extended Data Fig. 1b-c).

The same AI-boosted molecular MRI method was then translated to a clinical human MRI scanner, using the same encoding-decoding procedure (Fig. 1) and only minimal modifications to the pulse sequence to minimize tissue RF power deposition (see the Methods section and Extended Data Table 1 for additional details). A healthy volunteer was recruited and imaged at 3T, following Institutional Review Board (IRB) approval and informed consent. The resulting molecular maps (Fig. 4) yielded semi-solid proton volume fractions of 9.4±3.0% and 4.2±4.4% for white matter (WM) and gray matter (GM) regions, respectively. The elevated semi-solid proton volume fraction observed in WM compared to GM is consistent with the higher lipid/myelin content of WM. The resulting semi-solid exchange rates were WM: 14.0±6.9 Hz; GM: 35.1±15.4 Hz. Since no difference in pH is expected between WM and GM, the difference in semi-solid exchange rate observed for WM and GM again indicates the sensitivity of the semi-solid base catalyzed exchange rate constant to the lipid composition as also observed in the mouse tumor model (Fig. 2d). Both semi-solid exchange parameter values were in good agreement with previous studies (volume fractions: 13.9±2.8% and 5.0±0.5%, exchange rates: 23±4 and 40±1 Hz, for the WM and GM, respectively)^9^, despite the substantial variance existing in the literature (Extended Data Table 2). The measured amide proton WM/GM exchange rates (42.3±2.9 Hz / 34.6±9.5 Hz) were in good agreement with previous Water Exchange Spectroscopy (WEX) measurements in rat models (28.6±7.4 Hz)^29^. All data are presented as mean ± standard deviation.

**Figure 4:**
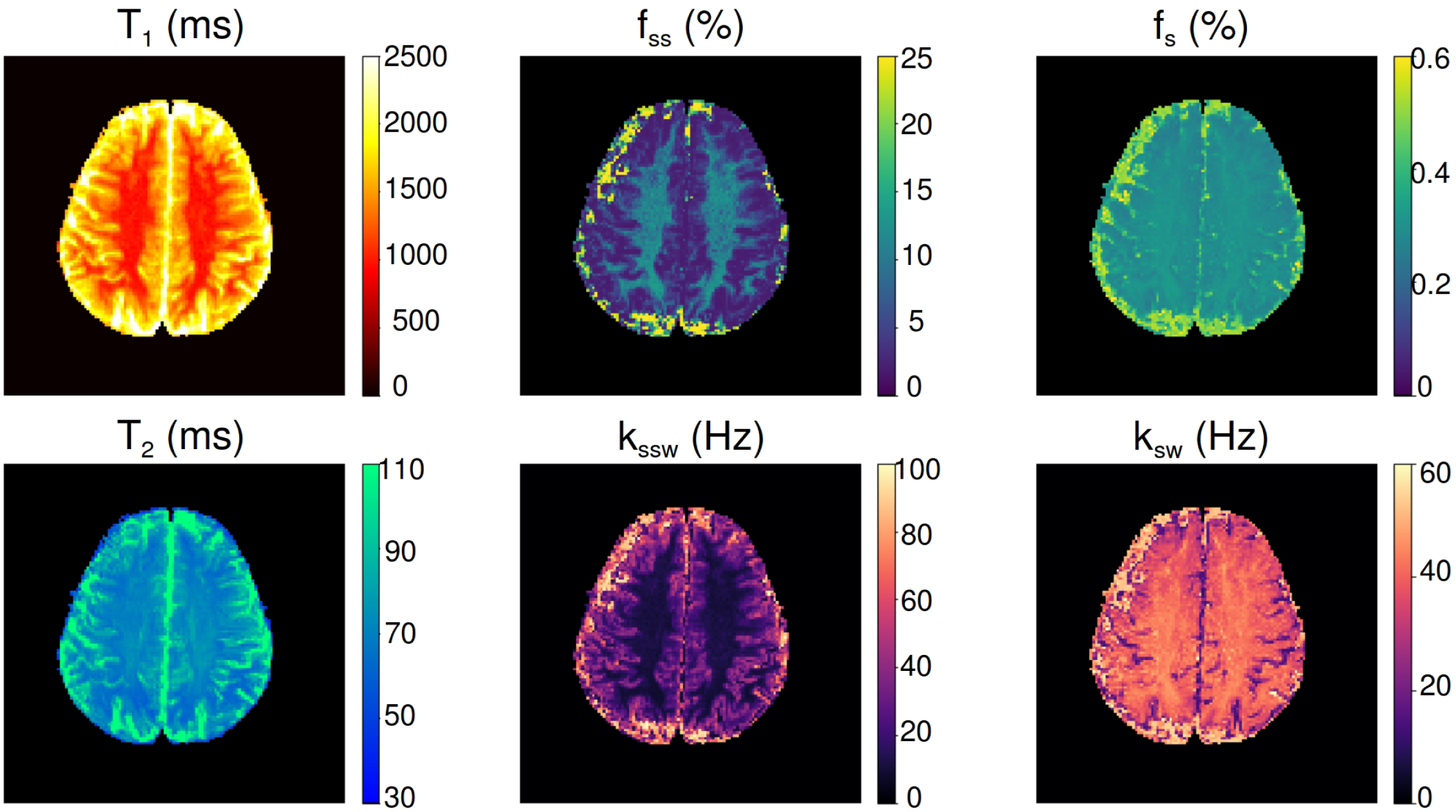
Clinical translation of the AI-boosted molecular MRI method and its evaluation on a healthy volunteer at 3T. The resulting white/gray-matter semi-solid volume fractions (9.4±3.0% / 4.2±4.4%) and exchange-rates (14.0±6.9 Hz / 35.1±15.4 Hz) were in good agreement with the literature (see Extended Data Table 2). The white/gray-matter amide proton exchange-rates (42.3±2.9 Hz / 34.6±9.5 Hz) were in good agreement with previous Water Exchange Spectroscopy (WEX) measurements in rat models^29^.

## Discussion

Apoptosis is considered an early predictor of cancer therapy outcome, as it manifests prior to any visible reduction in tumor volume^30, 31^. This has motivated the pursuit of an imaging method capable of apoptosis detection^32^. Previous methods developed for the detection of apoptosis relied on pH sensitive dual emission fluorescent probes^26^, caspase-3 targeted optical imaging probes^33, 34^, and radio-labeled Annexin V^35^ and duramycin^8^ positron emission tomography (PET) and single-photon emission computed tomography (SPECT) imaging probes that respectively target phosphatidylserine or phosphatidylethanolamine expressed on the cell surface of apoptotic cells. Additional methods relied on the changes in the endogenous lipid proton MR-spectroscopy^30^(MRS) or high frequency ultrasound signals^36^. However, PET and SPECT require the use of ionizing radiation and exogenous probes while optical and ultrasound based methods suffer from a limited tissue penetration ability and are unsuitable for clinical neurological applications. In contrast, MRI provides a safe and clinically relevant alternative. The intuitive method of choice for MR-based apoptosis molecular imaging is MRS^30^. However, MRS is limited by a very poor spatial resolution and exceedingly long scan times due to the low sensitivity^37^. Diffusion weighted MR imaging was shown to be correlated with apoptosis processes^31^. However, it has low sensitivity and may result in false-positive apoptosis indications since numerous other biological/pathological processes induce diffusion changes^7^. Exogenous contrast agents, such as gadolinium or superparamagnetic iron oxides (SPIO) can be administered for MR T_1_/T_2_-based apoptosis detection, after labeling the contrast agent with an appropriate targeting probe^6^. However, due to the difficulty in delivering large targeted contrast agents across the blood-brain barrier and recent concerns of the risk of adverse events when using metal-based probes for contrast enhancement, an endogenous-contrast-based method would constitute a favorable alternative.

CEST-MRI of endogenous amide protons has been extensively explored for pH-weighted imaging; hence, it constitutes a potential tool for apoptosis detection. The CEST pH imaging ability was initially demonstrated in acute stroke rodent models^10^ and later translated into clinical scanners and human subjects^38^. However, such studies typically use the magnetization transfer ratio asymmetry (MTR_*asym*_) analysis metric, which is affected by the proton exchange rate and volume fraction, by aliphatic proton pools (rNOE), by the water T_1_ and T_2_ relaxation times, and the saturation pulse properties. As a result, pathology-related changes in the water pool T_1_ and T_2_ and semi-solid and aliphatic proton pool properties may challenge the correct interpretation of a qualitative CEST-weighted image^39^ (Extended Data Fig. 2a). Moreover, although the various contributions of different proton pools to the CEST contrast can be separated-out using the previously suggested Lorentzian fitting approach^40^, the output, in this case, is still a single pool-weighted image for each molecular compound that cannot fully distinguish between pH changes and protein concentration or magnetic relaxation effects (Extended Data Fig. 2b-f). Therefore, in the presence of competing molecular mechanisms the resulting CEST-weighted images will not necessarily be correlated with changes in either pH or protein content. This is demonstrated in the conventional Lorentzian fitted parameters obtained for the virotherapy-treated mice, shown in Extended Data Fig. 3a. Here, despite the increase in exchange rate and decrease in proton volume fraction observed in the tumor for the quantitative CEST fingerprinting maps (Fig. 2a), no significant difference in amide proton CEST contrast is observed between contralateral and tumor tissue. This is attributed to the fact that the CEST contrast is proportional to the product of the proton exchange rate and concentration. In these challenging cases, a quantitative approach such as MR-fingerprinting^23^ is very attractive.

Nevertheless, a straight-forward implementation of a single CEST MR-fingerprinting encoding scheme^41^, and the traditional correlation-based reconstruction provide a very poor estimation of the molecular properties in in-vivo disease cases, such as cancer (Extended Data Fig. 2g-l). This stems from the highly multi-dimensional parameter-space involved and the simultaneous parameter variations in multiple molecular pools. The sequential architecture of the AI-boosted CEST MR-fingerprinting approach proposed here, overcomes all of the above challenges. Specifically, our approach first uses a semi-solid macromolecule selective encoding to properly isolate and quantify the properties of this pool (Fig. 1b). Only then, using a much smaller parameter-space, are the amide proton properties mapped with the use of an amide-oriented encoding schedule (Fig. 1b). The ability of the sequential deep networks to computationally manage large numbers of parameters allows us to properly include the effects of the water pool relaxation properties and magnetic field inhomogeneity as inputs to both neural networks. Extended Data Fig. 2m-r further demonstrates that sequentially nailing-down each pool parameters before classifying the next pool is indeed an essential strategy for overcoming this highly multidimensional challenge. Moreover, the use of neural-networks for image reconstruction allows for continuous parameter classification, instead of the discrete set of values obtained when performing traditional correlation-based matching where the matching dictionary contains only certain discrete parameter values. Finally, the image reconstruction time is 88,085 times shorter for the proposed method compared to standard MR-fingerprinting (94 ms instead of 2.3 hours, for a 128 × 128 pixel image). Our use of neural networks in this fingerprinting approach differs from the use of neural network based non-linear regression methods which were recently used for the extraction of proton pool Lorentzian parameters^42^ and quantification of phosphocreatine in leg muscle^43^ from conventional CEST Z-spectra. In particular, the fingerprinting method has previously been shown to have significantly improved parameter discrimination compared to CEST Z-spectra^41^. The combination of the fingerprinting method with the sequential deep networks is critical for characterizing the much more complicated tumor tissue pathology, where a very large number of different molecular parameters are all changing and must be accounted for in the model.

In terms of the image acquisition time, it is noted that the requirement of static magnetic-field (B_0_) and water-pool relaxation parameter maps (Fig. 1c) as network inputs prolongs the total scan-time. However, all 3 maps can be obtained in approximately 30 seconds using the previously established water-pool T_1_ and T_2_ MR-fingerprinting method^23^. Thus, with a 3.5 minute acquisition time for the amide and semi-solid proton pools, the total clinical acquisition time is less than 5 minutes. Notably, it is highly desirable to convert the single-slice method implemented here into a 3D protocol, providing whole brain coverage. This could be pursued by combining the CEST MR-fingerprinting approach with fast volumetric acquisition protocols, such as 3D-snapshot CEST^44^, multiband simultaneous multi-slice EPI^45^, or multi-inversion 3D EPI^46^.

The non-invasive apoptosis monitoring ability presented here could be expanded and become beneficial for a variety of additional clinical scenarios, characterized by an irregular apoptosis-level. This includes liver disease^47^, transplant rejection imaging^48^, Alzheimers, Parkinsons and Huntington disease^49^. More generally, although the main studied application for the AI-boosted molecular MRI was oncolytic virotherapy, the method is directly applicable to any semi-solid macromolecule and amide proton CEST imaging application. This includes pH imaging for stroke detection and ischemic penumbra characterization^10^, differentiation of ischemia from hemorrhage^50^, cancer grading^11^, detection of biological therapeutics engineered with CEST reporter genes^5, 51, 52^, differentiation of radiation necrosis and tumor progression^13^, multiple sclerosis lesion detection and evaluation^53^, and neurodegenerative disease characterization^54^. Furthermore, future work could optimize and modify the encoding scheme (Fig. 1b) so that additional metabolites and molecular information could be quantitated (e.g., creatine, glutamate, and glucose), opening the door for a plethora of new and exciting molecular insights.

## Conclusion

The AI-boosted molecular MRI method successfully and rapidly quantified proton chemical exchange rate and protein concentration based changes, serving as important biomarkers for the detection of oncolytic virotherapy induced cell apoptosis.

## Acknowledgments

The work was supported by the US National Institutes of Health Grants R01-CA203873, P41-RR14075, 1S10RR023401, 1S10RR019307, and 1S10RR023043 as well as by 1R01NS110942-01A1 and P01 CA163205 (EAC and HN). The Brigham and Womens Small Animal Imaging Laboratory (SAIL) was funded by a G20 Grant (1G20RR031051-01) as part of the American Recovery and Reinvestment Act as a part of the construction of the Brigham and Womens MRI Research Center (BWMRC). The 7T Bruker Small Bore Animal Magnet was partially funded by an S10 Grant (1S10OD010705-01) through the National Institutes of Health. K.H was supported by the German Research Foundation (DFG, grant ZA 814/21). This project has received funding from the European Unions Horizon 2020 Research and Innovation Programme under the Marie Sklodowska-Curie grant agreement No. 836752 (OncoViroMRI).

## Author contributions

O.P., O.C., M.S.R., and C.T.F. conceptualized the deep-learning reconstruction architecture. O.P., K.H., M.Z., O.C., and C.T.F. implemented and tested various aspects of the technical framework. H.I., H.N., E.A.C., O.P., and C.T.F. contributed to preclinical experiment design. H.I. performed tumor implantations and virus inoculations. O.P., C.T.F., and H.I. performed the preclinical imaging. N.S. performed the histology and immunohistochemistry studies. K.H., M.Z., O.P., and C.T.F. developed the clinical imaging scheme. K.H. and M.Z. performed the clinical imaging. O.P., H.I., K.H., N.S., H.N., M.Z., E.A.C., O.C., M.S.R., and C.T.F. wrote and/or substantially revised the manuscript.

## Methods

### In vivo preclinical MRI acquisition

The mouse imaging study was conducted using a 7T pre-clinical MRI scanner (Bruker Biospin, Ettlingen, Germany). Two chemical exchange saturation transfer (CEST) MR-fingerprinting acquisition protocols were employed sequentially (105s each, Fig. 1a-b), designed for encoding the semi-solid macromolecule (denoted as MT, magnetization transfer) and amide information into unique trajectories. The exchangeable amide proton signals of endogenous mobile proteins has a chemical shift of 3.5 ppm with respect to water and a relatively narrow resonance linewidth due to the rapid molecular motion. In contrast, the exchangeable protons on lipids and large macromolecules have a chemical shift of approximately −2.5 ppm from water and a very broad resonance linewidth due to the slow molecular motion. The first protocol was aimed for magnetic labeling the MT pool, varying the saturation pulse frequency offset between 6-14 ppm (to avoid amide/amine/aliphatic nuclear Overhauser effect (NOE) contributions) and the saturation pulse power between 0-4 *µ*T. The second protocol was aimed for magnetic labeling the amide pool (in the presence of MT), using the same saturation pulse power schedule but with a fixed saturation pulse frequency offset at 3.5 ppm. Both protocols had repetition-time/echotime (TR/TE) = 3500/20 ms, a flip angle of 90, a continuous saturation pulse of 2500 ms, and a spin-echo echo-planar-imaging (SE-EPI) readout. T_1_ maps were acquired using the variable repetition-time rapid acquisition with relaxation enhancement (RARE) protocol, with TR = 200, 400, 800, 1500, 3000, and 5500 ms, TE = 7 ms, RARE factor = 2. T_2_ maps were acquired using the multi-echo spin-echo protocol, TR = 2000 ms, 25 TE values between 8-200 ms. Static magnetic field B_0_ maps were acquired using the water saturation shift referencing (WASSR) protocol^55^, employing a saturation pulse power of 0.3 *µ*T, TR/TE = 8000/20 ms, flip-angle = 90, saturation duration = 3000 ms, and a saturation pulse frequency offset varying between −1 to 1 ppm in 0.1 ppm increments. A traditional full Z-spectrum CEST scan was performed for comparison, using a SE-EPI sequence with TR/TE = 8000/20 ms, flip-angle = 90, and pre-saturation pulses of 0.7 *µ*T and 3000 ms, at −7 to 7 ppm frequency offsets with 0.25 ppm increments and a no-saturation reference image. The field of view (19 mm × 19 mm × 1 mm) and image resolution (297 × 297 × 1000 *µ*m^3^) were identical for all scans besides a high-resolution (148 × 148 × 1000 *µ*m^3^) T_2_-weighted scan (TR/TE = 2000/60 ms), taken as reference.

### CEST MR-fingerprinting dictionary generation

Dictionaries of CEST signals were generated, simulating the expected trajectories for more than 70 million tissue parameter combinations, as a response to the two molecular-information-encoding acquisition protocols (Fig. 1a-b). The simulations were carried out using a numerical solution of the Bloch-McConnell equations, implemented in MATLAB R2018a (The MathWorks, Natick, MA) and C++ ^41, 56^. Generating all dictionaries used in this work took a total of 62.5 hours, using a computer cluster employing 56 CPUs. Detailed dictionary properties can be found in Extended Data Table 1.

### Quantitative image decoding using deep reconstruction networks

To avoid the exceedingly long dictionary matching-time required for conventional correlation-based MR-fingerprinting (e.g., 2.31 hours for reconstructing a single 128 × 128 pixels image-set out of >70M dictionary entries, using a 12 GiB Intel Xeon E5607 CPU equipped Linux desktop computer) and to improve the multi-parameter reconstruction ability (Extended Data Fig. 2g-r), image decoding was performed using a series of two deep reconstruction networks (DRONEs)^57^. Each neural-network was comprised of 4-layers, including 300 × 300 neurons in the two hidden layers (Fig. 1c). A rectified linear unit (ReLU) and a sigmoid were used as the hidden and output activation functions, respectively. Network training was performed using the synthesized dictionary data, with the trajectories normalized to zero mean and unit standard deviation along the temporal axis^58^. The adaptive moment estimation (ADAM) optimizer^59^ was used with a learning rate = 0.0001, minibatch size = 256, and the mean-squared-error defined as the loss-function. To avoid over-fitting, 10% of each dictionary (Extended Data Table 1) was excluded from training and was used to assess when to stop the training (early stopping). To promote robust learning, white Gaussian noise (signal-to-noise ratio (SNR) = 20 dB) was injected into the dictionaries^60^. At the reconstruction step, the pixel-wise signal trajectories from the 30 images acquired using the MT-specific MR-fingerprinting schedule were normalized along the temporal axis^58^ and input to the first DRONE, together with the pixel-wise water T_1_, T_2_ and B_0_ values (normalized by subtracting the mean and dividing by the standard deviation of the training dictionary parameters). The two MT exchange parameter output maps, together with the water pool T_1_ and T_2_ relaxation and B_0_ parameter maps were then input into the second DRONE, together with the pixel-wise signal trajectories from the 30 images acquired using the amide-pool MR-fingerprinting schedule (normalized similarly to the MT schedule images). The neural-networks were implemented in Python 3.6 using TensorFlow 1.4.1, on a desktop computer equipped with an Intel Xeon E5607 CPU and an NVIDIA TITAN Xp GPU.

### Animal model

Six-to-8-week-old female athymic mice (BALB/c, nu/nu) were purchased from Envigo. All animal experiments were performed in accordance with protocols approved by the Brigham and Women’s Hospital Institutional Animal Care and Use Committee (IACUC). 100,000 cells of U87ΔEGFR were implanted stereotactically with a Kopf stereotactic frame (David Kopf Instruments) in the right frontal lobe of 16 mice (ventral 3.0 mm, rostral 0.5 mm, and right lateral 2.0 mm from bregma). Imaging was performed at 8-11 days post implantation, using a 7T pre-clinical MRI (Bruker Biospin, Ettlingen, Germany). Next, a herpes simplex virus type I-derived oncolytic virus, NG34^61^, was inoculated intratumorally for 12 mice (the others served as control), and the imaging was repeated 48 and 72 hours later. The mice were anesthetized using 0.5 to 2% Isoflurane during the imaging on a dedicated heated cradle, and the respiration rate was continuously monitored using a mechanical sensor. Following the last imaging time-point, the mice were euthanized using CO_2_ inhalation and the brain-tissue extracted, formalin-fixed and paraffin-embedded for histology and immunohistochemistry. An additional tumor-bearing mouse (without treatment) was used for demonstrating the differences between the proposed and previously suggested molecular CEST MRI methods (Extended Data Fig. 2). Four mice died before the planned termination point (two between the baseline and 48 hours scan and two between the 48 hours and 72 hours scan).

### Statistical analysis

Comparative analyses were carried out using one-way ANOVA with posthoc Tukey’s multiple comparisons test, using Prism 6 (GraphPad Software, Inc, La Jolla, CA). Differences were considered significant at P<0.05. Box plots (Fig. 2d, Extended Data Fig. 3) represent the median, interquartile range, Tukey style whiskers, and outliers. Column scatter plots (Extended Data Fig. 1) include horizontal and vertical lines representing the group mean and standard deviation, respectively. The following two exclusion criteria were imposed: unsuccessful tumor implantation (occurred in 1 mouse out of 16); corrupted image data (occurred in 2 image-sets out of 39, potentially due to RF coil/transmitter error).

### Immunohistochemistry

Mouse brain tissues were fixed with 10% neutralization buffer and embedded in paraffin by Servicebio Inc (Woburn, MA). The embedded samples were sectioned and processed sequentially with xylene, ethanol, distilled water to attain deparaffinization and rehydration. The following procedures were performed separately for each staining.

#### Immunohistochemistry

Histology slides were kept in citrate buffer (pH6) heated to sub-boiling temperature for 20 minutes and cooled at room temperature for 30 minutes. After treating the slides with 3% hydrogen peroxide in distilled water to lower intrinsic peroxidase activity, the slides were incubated with 2% normal goat serum/20 mM Tris-Buffered Saline, 0.05 % Tween-20 (TBST) to block unspecific antibody binding. Slides were incubated with primary antibodies against HSV 1 (B0114, Dako; 1:100 dilution) or cleaved caspase 3 (9579, Cell Signaling Technology; 1:150) diluted as recommended by the manufacturer with TBST. The MACH4 Universal HRP-Polymer (M4U534, Biocare Medical) and Metal Enhanced DAB Substrate Kit (34065, Thermo Fisher Scientific) was used to induce chromogenic reaction for detection. Additionally, the tissue was counterstained with Hematoxylin.

#### HE stain

The slides were stained with Mayer’s Hematoxylin (MHS, Sigma-Aldrich) and Eosin (HT110, Sigma-Aldrich).

#### Coomassie stain

A Coomassie stain was performed for the detection of protein concentration as described by Ray et al.^62^.

Stained slides were imaged with the Nikon Ti Eclipse microscope system (Nikon, Minato-ku, Japan) and captured by NIS 5.11.01 software (Nikon, Minato-ku, Japan).

### Clinical translation

The same imaging approach implemented in the animal study was translated for clinical scanners and human subjects, with minimal modifications, as mandated by the difference in hardware and specific absorption rate (SAR) restrictions. Specifically, the continuous-wave saturation pulse was replaced by a train of off-resonant spin-lock saturation pulses (13 × 100 ms, 50% duty-cycle^63^), and the read-out was done using gradient-echo (GRE) EPI. The MT and amide specific MR-fingerprinting protocols were realized using the hardware-independent open-source pulseq framework^64^ in MATLAB, with the same saturation pulse power and frequency offsets used in the preclinical study (Fig. 1b), and were played out by an interpreter on the scanner. T_1_ and T_2_ mapping were performed using saturation recovery and a series of five single-echo spin-echo sequences with different TE, respectively. B_0_ maps were acquired using the WASABI method^65^. A healthy volunteer (27 year old male) was recruited following the University of Tübingen IRB approval and informed consent and imaged at 3T (Siemens Healthineers, Germany). All images had the same resolution of 1.72 × 1.72 × 10 mm^3^. The decoding of the quantitative molecular information was performed similarly to the preclinical study, using deep reconstruction networks trained with dictionaries simulated using the clinical acquisition protocol (Extended Data Table 1).

### Image analysis

Data analysis was performed using MATLAB R2018a and Python 3.6 custom written scripts, based on previously published routines, as described below. T_1_ and T_2_ map reconstructions were performed using exponential fitting. Conventional CEST images were corrected for B_0_ in-homogeneity using the WASSR method^55, 66^, followed by cubic spline smoothing^67, 68^. The magnetization transfer ratio asymmetry (Extended Data Fig. 2a) was calculated using: MTRasym = (S^−Δ*ω*^ – S^+Δ*ω*^) / S_0_, where S^±Δ*ω*^ is the signal measured with saturation at offset ± 3.5 ppm and S_0_ is the unsaturated signal. Semi-quantitative mapping of the CEST molecular compound amplitudes was performed using a 5-pool (water, MT, amide, NOE, and amine) Lorentzian fitting model (Extended Data Fig. 3), with the starting point and boundaries described in^40^. An image down-sampling expedited adaptive least-squares (IDEAL) fitting approach was then implemented (Extended Data Fig. 2b-f), as described in^69^. CEST MR-fingerprinting with conventional dictionary matching (Extended Data Fig. 2g-l) was performed by calculating the dot-product after 2-norm normalization of each amide encoded image pixel trajectory with all relevant dictionary entries (Extended Data Table 1)^41, 56^. Mouse tumor regions of interest (ROIs) (Fig. 2, Extended Data Fig. 1) were manually delineated based on the T_1_ and T_2_ maps. The contralateral ROIs were automatically obtained by symmetrically reflecting the tumor ROIs. Suspected apoptotic ROIs were delineated based on decreased amide proton exchange rate. If a suspected apoptotic region existed, its area was excluded from the tumor ROI. All delineations were performed by a single observer blinded to the histology. Human white matter and gray matter ROIs were automatically segmented based on the T_1_-map and literature T_1_ values at 3T^9^, allowing a margin of three standard deviations from the mean.

### Data availability

The datasets generated and analysed during the current study are available from the corresponding author upon request.

### Code Availability

Source code is available from the corresponding author upon request.

---

**Extended Data Fig. 1:**
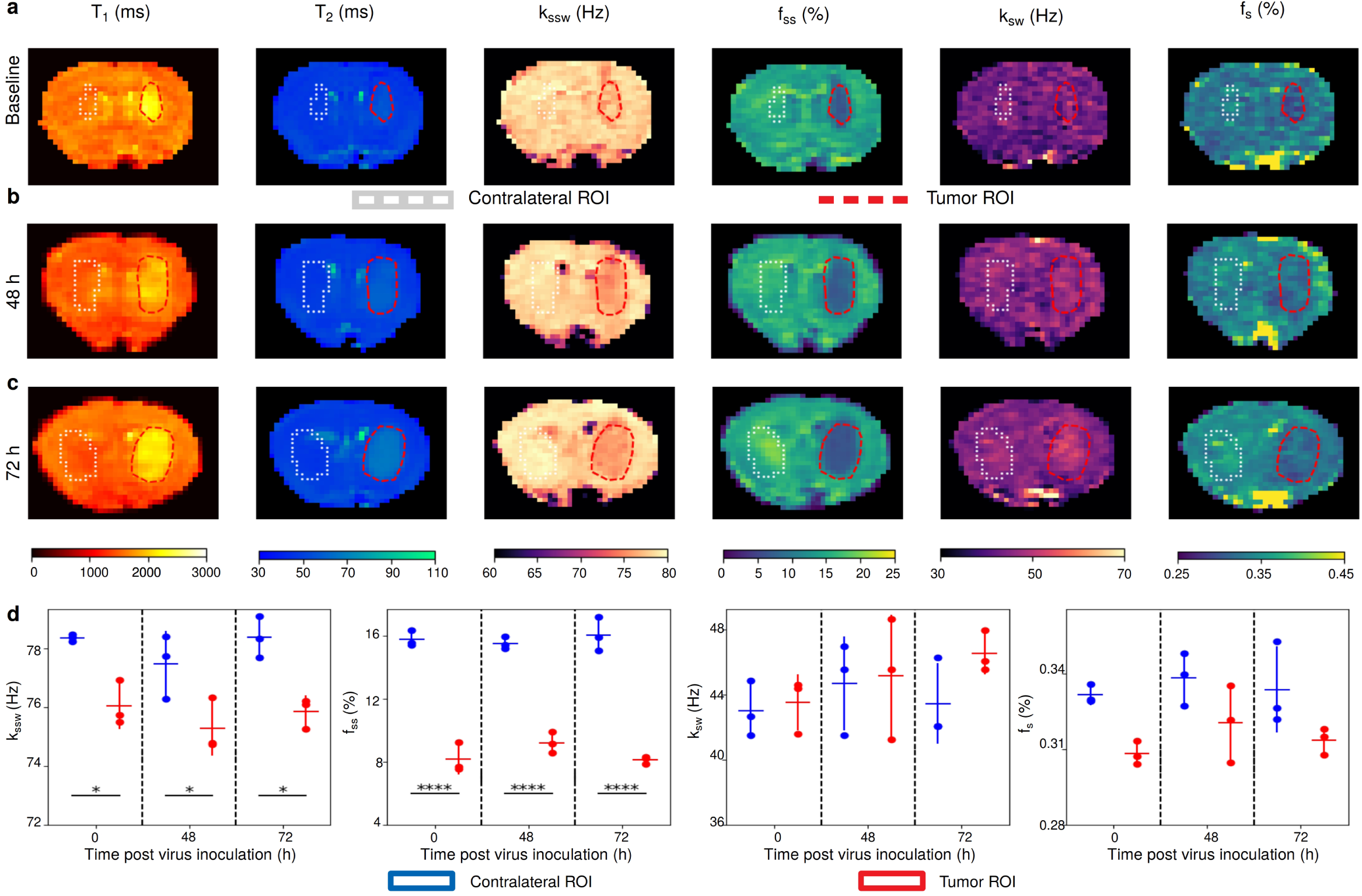
Quantitative molecular images of a control (tumor-bearing, untreated) mouse. a. Baseline. The tumor semi-solid (f_*ss*_) and amide (f_*s*_) proton concentrations were decreased, consistent with increased edema. The tumor amide proton exchange-rate (k_*sw*_) was increased, indicative of increased intracellular pH. Forty-eight (b) and 72 (c) hours following the baseline scan, no apoptosis was detected, as expected. d. Column scatter plot display of the group exchange parameters. One-way analysis of variance (ANOVA): F(5, 12)=9.598, 100.8, 3.804, 0.8968; P=0.0007, P<0.0001, P=0.0269, P=0.5136, for semi-solid exchange rate (k_*ssw*_), f_*ss*_, f_*s*_, and k_*sw*_ respectively. Horizontal and vertical color lines represent the group mean and standard deviation, respectively. *P<0.05; ****P<0.0001 (Tukeys multiple comparison test, n = 3).

**Extended Data Fig. 2:**
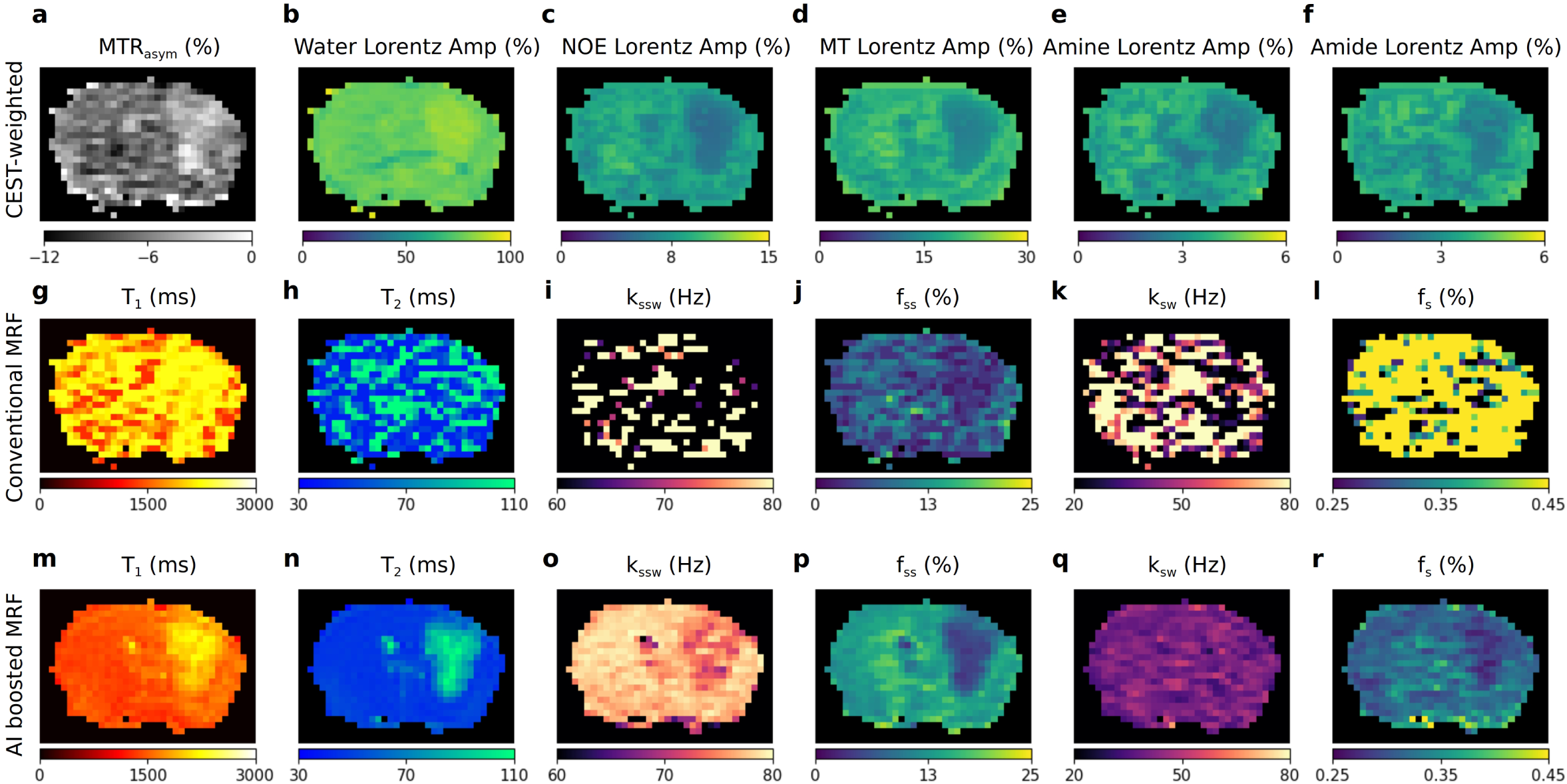
Comparison of chemical exchange saturation transfer (CEST) MRI methods on a brain tumor-bearing mouse. a. Typical amide proton CEST-weighted MTR_*asym*_ image. The contrast is affected by a multitude of molecular compounds and protocol-specific acquisition parameters. b-f. Lorentzian fitting of the CEST Z-spectrum provides a semi-quantitative map for each of the modeled compounds (water, aliphatic NOE, semisolid-MT, amine, and amide). Each map represents the combined effect of the exchange rate, volume fraction, and water T_1_ and T_2_. g-l. CEST MR-fingerprinting with conventional dictionary matching. Note the resulting poor SNR. m-r. AI boosted CEST-MRF, which successfully decouples and quantifies the exchange parameters for each exchangeable proton pool.

**Extended Data Fig. 3:**
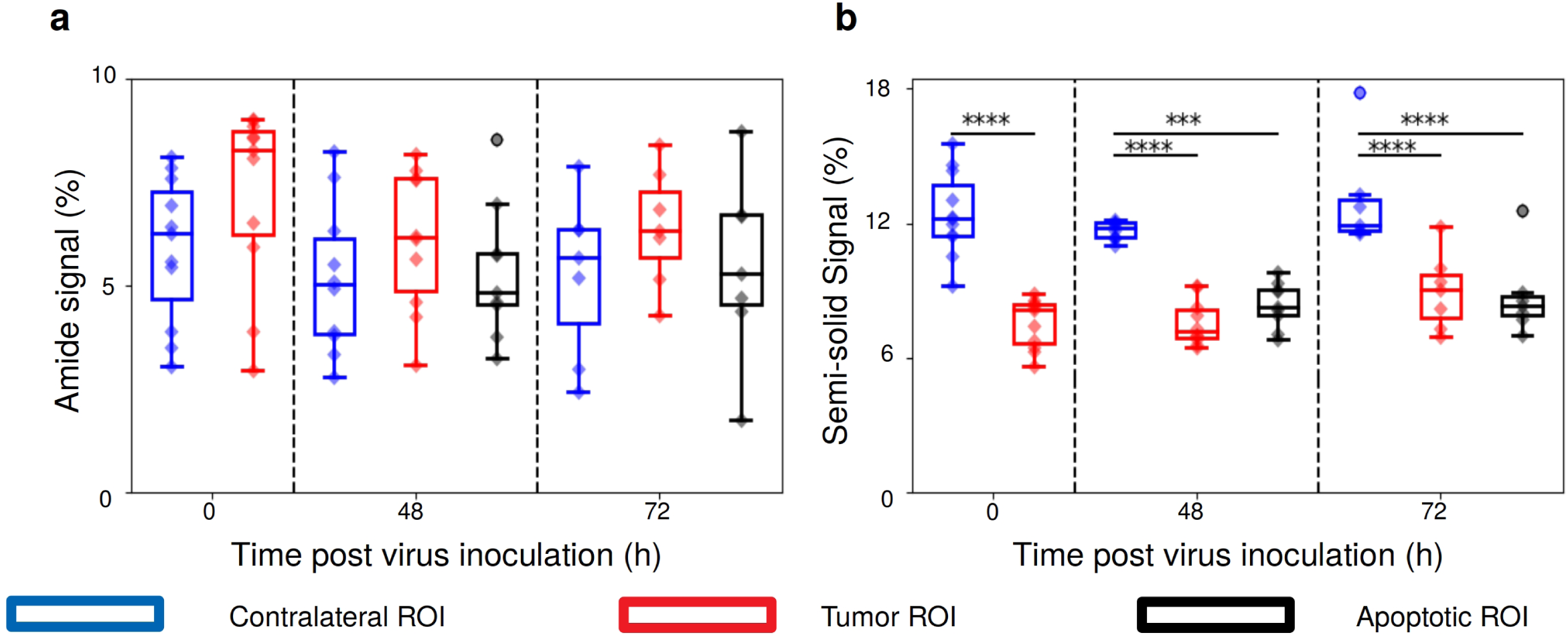
Quantitative group comparison for traditional CEST Z-spectrum Lorentzian fitting for the virotherapy mice. a. Amide proton signal. No significant difference in amide CEST signal is observed between contralateral and tumor tissue, ANOVA (F(7, 64)=1.445, P=0.2031). This is due to the opposite trends of the contrast affecting mechanisms (increased amide proton exchange rate and decreased amide proton volume fraction, Fig. 2a). b. Semi-solid proton signal. Significant signal decrease was measured between the contralateral and tumor/apoptotic regions, in agreement with the simultaneous decrease observed in both the semi-solid proton volume fraction and exchange rate (Fig. 2d). ANOVA (F(7, 64)=21.90, P<0.0001). Asterisks represent the P-value for Tukeys multiple comparison test: ***P<0.001; ****P<0.0001, for comparing the different ROI signals for baseline (n=11 all regions), 48 hours (n=10 contralateral and tumor; n=9, apoptotic), and 72 hours (n=7, all regions) post inoculation. The box plots describe the median, Tukey style whiskers, and outliers.

**Extended Data Table 1.**
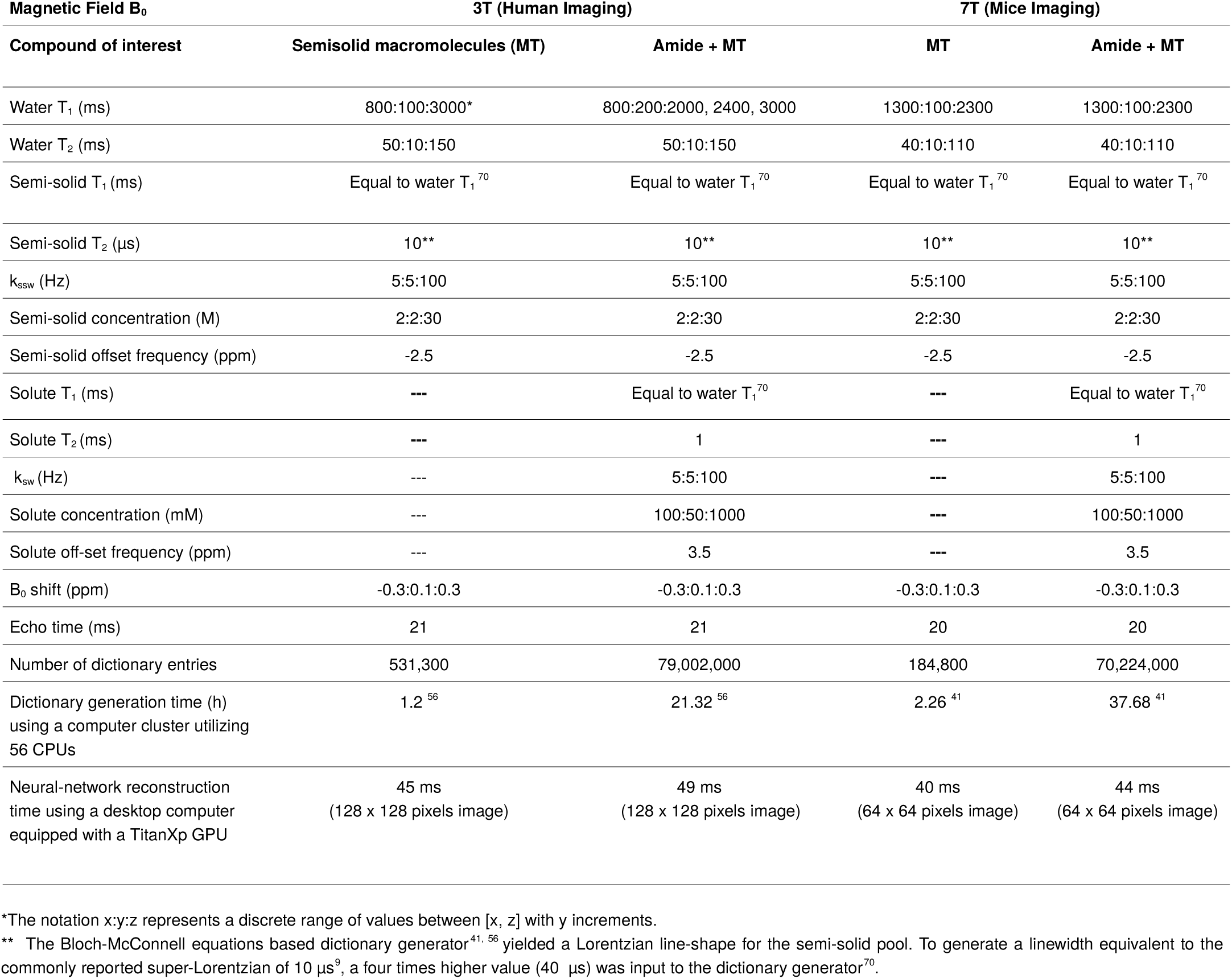
Detailed properties of the MR-fingerprinting dictionaries used for training the deep reconstruction networks.

| Magnetic Field B<sub>0</sub> | 3T (Human Imaging) |  | 7T (Mice Imaging) |  |
| --- | --- | --- | --- | --- |
|  | Semisolid macromolecules (MT) | Amide + MT | MT | Amide + MT |
| Water T<sub>1</sub> (ms) | 800:100:3000* | 800:200:2000, 2400, 3000 | 1300:100:2300 | 1300:100:2300 |
| Water T<sub>2</sub> (ms) | 50:10:150 | 50:10:150 | 40:10:110 | 40:10:110 |
| Semi-solid T<sub>1</sub> (ms) | Equal to water T<sub>1</sub><sup>70</sup> | Equal to water T<sub>1</sub><sup>70</sup> | Equal to water T<sub>1</sub><sup>70</sup> | Equal to water T<sub>1</sub><sup>70</sup> |
| Semi-solid T<sub>2</sub> (μs) | 10** | 10** | 10** | 10** |
| k<sub>ssw</sub> (Hz) | 5:5:100 | 5:5:100 | 5:5:100 | 5:5:100 |
| Semi-solid concentration (M) | 2:2:30 | 2:2:30 | 2:2:30 | 2:2:30 |
| Semi-solid offset frequency (ppm) | -2.5 | -2.5 | -2.5 | -2.5 |
| Solute T<sub>1</sub> (ms) | --- | Equal to water T<sub>1</sub><sup>70</sup> | --- | Equal to water T<sub>1</sub><sup>70</sup> |
| Solute T<sub>2</sub> (ms) | --- | 1 | --- | 1 |
| k<sub>sw</sub> (Hz) | --- | 5:5:100 | --- | 5:5:100 |
| Solute concentration (mM) | --- | 100:50:1000 | --- | 100:50:1000 |
| Solute off-set frequency (ppm) | --- | 3.5 | --- | 3.5 |
| B<sub>0</sub> shift (ppm) | -0.3:0.1:0.3 | -0.3:0.1:0.3 | -0.3:0.1:0.3 | -0.3:0.1:0.3 |
| Echo time (ms) | 21 | 21 | 20 | 20 |
| Number of dictionary entries | 531,300 | 79,002,000 | 184,800 | 70,224,000 |
| Dictionary generation time (h)<br>using a computer cluster utilizing<br>56 CPUs | 1.2<sup>56</sup> | 21.32<sup>56</sup> | 2.26<sup>41</sup> | 37.68<sup>41</sup> |
| Neural-network reconstruction<br>time using a desktop computer<br>equipped with a TitanXp GPU | 45 ms<br>(128 x 128 pixels image) | 49 ms<br>(128 x 128 pixels image) | 40 ms<br>(64 x 64 pixels image) | 44 ms<br>(64 x 64 pixels image) |

\*The notation x:y:z represents a discrete range of values between [x, z] with y increments.
\*\* The Bloch-McConnell equations based dictionary generator<sup>41, 56</sup> yielded a Lorentzian line-shape for the semi-solid pool. To generate a linewidth equivalent to the commonly reported super-Lorentzian of 10 μs<sup>9</sup>, a four times higher value (40 μs) was input to the dictionary generator<sup>70</sup>.

**Extended Data Table 2.** Literature values of the brain white/gray matter semi-solid and amide proton volume fractions and exchange rates.

| Brain tissue | Semi-solid proton volume fraction (%) | Semi-solid proton exchange rate (Hz) | Amide proton volume fraction (%) | Amide proton exchange rate (Hz) | Reference |
| --- | --- | --- | --- | --- | --- |
| WM | 9.4 ±3.0 | 14.0±6.9 | 0.31 ± 0.02 | 42.3 ±2.9 | This work |
| GM | 4.2 ±4.4 | 35.1±15.4 | 0.32 ± 0.07 | 34.6 ±9.5 |  |
| WM | 13.9 ±2.8 | 23±4 |  |  | Stanisz et al., 2005<sup>9</sup> |
| GM | 5.0 ±0.5 | 40±1 |  |  |  |
| WM |  |  |  | 28.6±7.4* | van Zijl et al., 2003<sup>29</sup> |
| GM |  |  |  |  |  |
| WM | 8.9±0.3 |  | 0.21±0.03 |  | Geades et al., 2017<sup>71</sup> |
| GM | 4.4±0.4 |  | 0.20±0.02 |  |  |
| WM | 6.2±0.4 | 67.5±7.0 | 0.22±0.04 | 281.2±0.6 | Liu,et al., 2013<sup>72</sup> |
| GM | 3.4±0.4 | 63.5±4.5 | 0.25±0.05 | 281.9±0.9 |  |
| WM | 13.48±0.37 |  |  |  | Yarnykh, et al, 2015<sup>73</sup> |
| GM | 5.77±0.34 |  |  |  |  |
| WM | 12.1±0.4 | 21.05±0.5 |  |  | Samsanov, *et al*, 2012<sup>74</sup> |
| GM | 5.4±0.2 | 30.09±1.1 |  |  |  |
| WM | 11.2±0.7 | 29±4 | 0.19±0.02 | 162±16 | Heo et al., 2019<sup>75</sup> |
| GM | 6.3±0.7 | 40±5 | 0.24±0.02 | 365±19 |  |

WM = white matter; GM = gray matter
\*Measured using water exchange spectroscopy (WEX) in a rat cortex

